# Public human microbiome data dominated by highly developed countries

**DOI:** 10.1101/2021.09.02.458641

**Authors:** Richard J. Abdill, Elizabeth M. Adamowicz, Ran Blekhman

## Abstract

The importance of sampling from globally representative populations has been well established in human genomics. In human microbiome research, however, we lack a full understanding of the global distribution of sampling in research studies. This information is crucial to better understand global patterns of microbiome-associated diseases and to extend the health benefits of this research to all populations. Here, we analyze the country of origin of all 444,829 human microbiome samples that have been collected to date and are available from the world’s three largest genomic data repositories, including the Sequence Read Archive (SRA). We show that more than 71% of publicly available human microbiome samples with a known origin come from Europe, the United States, and Canada, including 46.8% from the United States alone, despite the country representing only 4.3% of the global population. We also find that central and southern Asia is the most underrepresented region: Countries such as India, Pakistan, and Bangladesh account for more than a quarter of the world population but make up only 1.8 percent of human microbiome samples. These results demonstrate a critical need to ensure more global representation of participants in microbiome studies.

## Background

A growing body of research shows the human microbiome has broad relevance to human health and disease. However, identifying the specific connections between the microbiome and human health requires a broad survey of both human populations and their most common health conditions. Even among healthy individuals, human microbiome composition varies between populations in ways that are still being uncovered: Geography and geographic relocation has been found to have an influence on microbiome composition (Yatsunenko et al. 2012; Vangay et al. 2018; Kaplan et al. 2019), as have host genetic variation and ethnicity (Goodrich et al. 2014; Blekhman et al. 2015; Brooks et al. 2018). Diet (Johnson et al. 2019), lifestyle (Clemente et al. 2015) and patterns in antibiotic use (Forslund et al. 2013) have all been linked to microbiome composition, with other studies considering the influence of locational factors such as pollution (Mutlu et al. 2018). Even within countries, interacting factors such as income, race, and education have critical impacts on health outcomes that could be mediated by the human microbiome (Amato et al. 2021). Some microbiome studies have specifically collected and compared data from global sites (Fragiadakis et al. 2019; Groussin et al. 2021), but large gaps and disparities still exist in which microbiomes are being studied on a global scale. The human microbiome has been linked to a growing number of social, medical and economic factors not directly related to host genetics, which reinforces the urgent need to evaluate the microbiomes of many populations (Amato et al. 2021; Ishaq et al. 2021).

Other genomics fields also have gaps in which populations are included in research. Genome-wide association studies (GWAS) have been primarily conducted in populations with European ancestry (Medina-Gomez et al. 2015; Gurdasani et al. 2019). As a result, polygenic risk scores from these studies have poorer accuracy when applied to non-European groups, limiting the possible benefits of this research—including personalized medicine, early disease screening, and risk prediction— to European-descended populations (De La Vega and Bustamante 2018; Peterson et al. 2019; Cai et al. 2021). There has been a concerted effort in genomics to include non-European individuals in GWAS studies, concurrent with calls to build research infrastructure and capacity globally (Gurdasani et al. 2019). It is likewise critical to identify underrepresented populations and locations in both genomics and microbiome research; otherwise, the benefits of host–microbiome research may only extend to a subset of the global population.

To investigate the geographic distribution of microbiome studies, we used metadata on all human microbiome datasets in the BioSample database, which includes metadata describing samples in the Sequence Read Archive (SRA), DNA Data Bank of Japan, and European Nucleotide Archive (Nakamura et al. 2013). Our data includes the country of origin and time of release for more than 444,000 samples, including both 16S amplicon sequencing and shotgun metagenomic sequencing, released over the last 11 years. These samples from the three largest genomic databases represent a large majority of all human microbiome samples that have been published.

## Results

We downloaded metadata for 444,829 human microbiome samples across 19 body sites and 2,592 studies. This data is available from the BioSample database maintained by the National Center for Biotechnology Information (NCBI), which includes metadata describing raw sequencing data deposited in multiple international repositories, including SRA (Barrett et al. 2012). While sample-level genomic sequencing data is uploaded to SRA, information such as subject age are saved separately to an entry in the BioSample database. BioSamples can be tagged with any number of “attributes,” including 485 standardized fields documented by NCBI (NCBI n.d.); we downloaded all attributes for all these samples. We used a Python script to load this metadata into a PostgreSQL database, where the information was aggregated using sample metadata such as country of origin and time of publication (see **Methods**).

As expected, we found the number of publicly available human microbiome samples has been increasing over time, from three microbiome samples in 2010 to 123,302 in 2020, the first year in which more than 100,000 human microbiome samples were released (**Supplementary Figure 1**). 2010 was the first year of the BioSample database, which all depositors must now use if they submit sequencing data to the Sequence Read Archive. The most common attribute in this subset of samples is the geographic origin of the sample (NCBI n.d.), which is available for 99.5% of samples (**Supplementary Table 1**). Using this attribute, we were able to determine the country of origin for 382,711 (86%) human microbiome samples (**Figure 1a**). We found that 178,960 samples (40.2%) are from the United States, almost five times more than any other country (**Table 1**). China has the next-most samples, with 36,162 (8.1%), followed by the United Kingdom, Denmark, Australia, and the Netherlands. China is the only Asian country in the top 14; the first South American country is Chile, in 16th place with 3616 samples (0.8%). Malawi is the first African country, in 19th place with 3052 samples (0.7%).

**Table 1.**
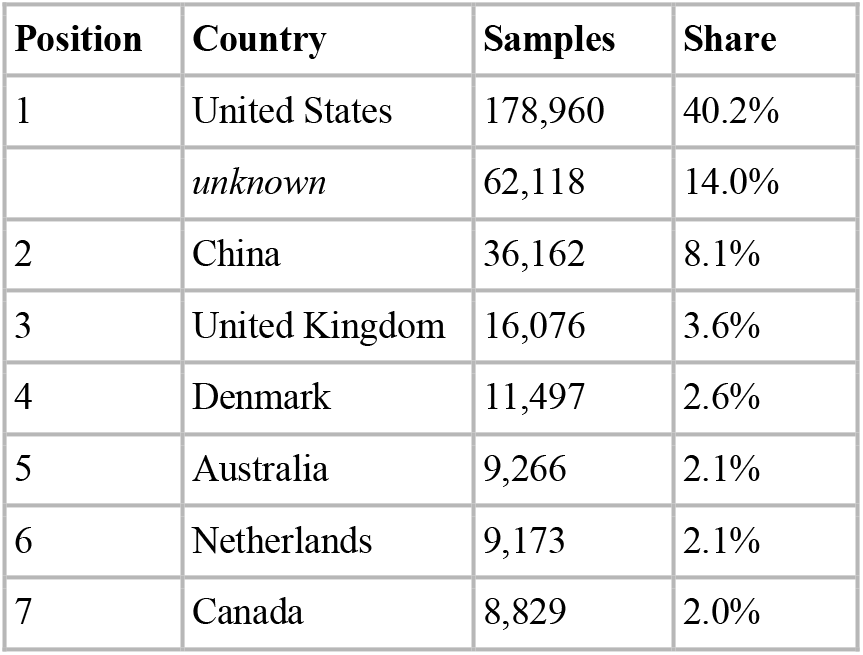
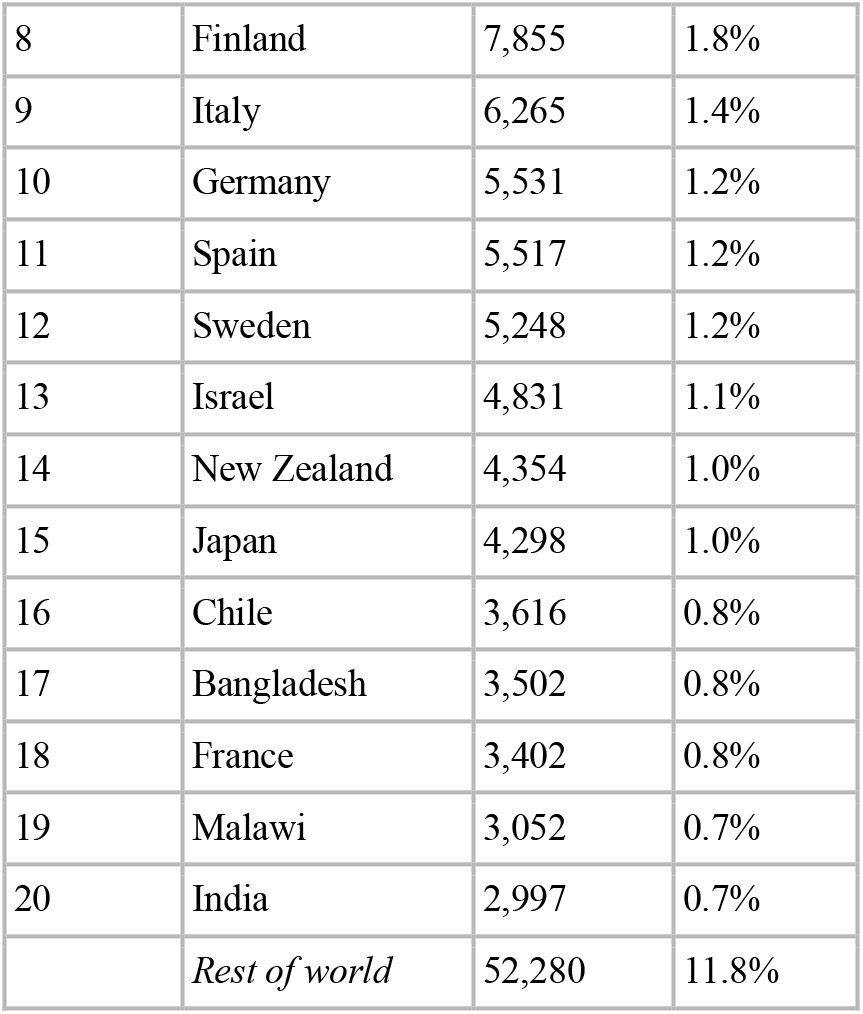
Samples per country.

**Figure 1.**
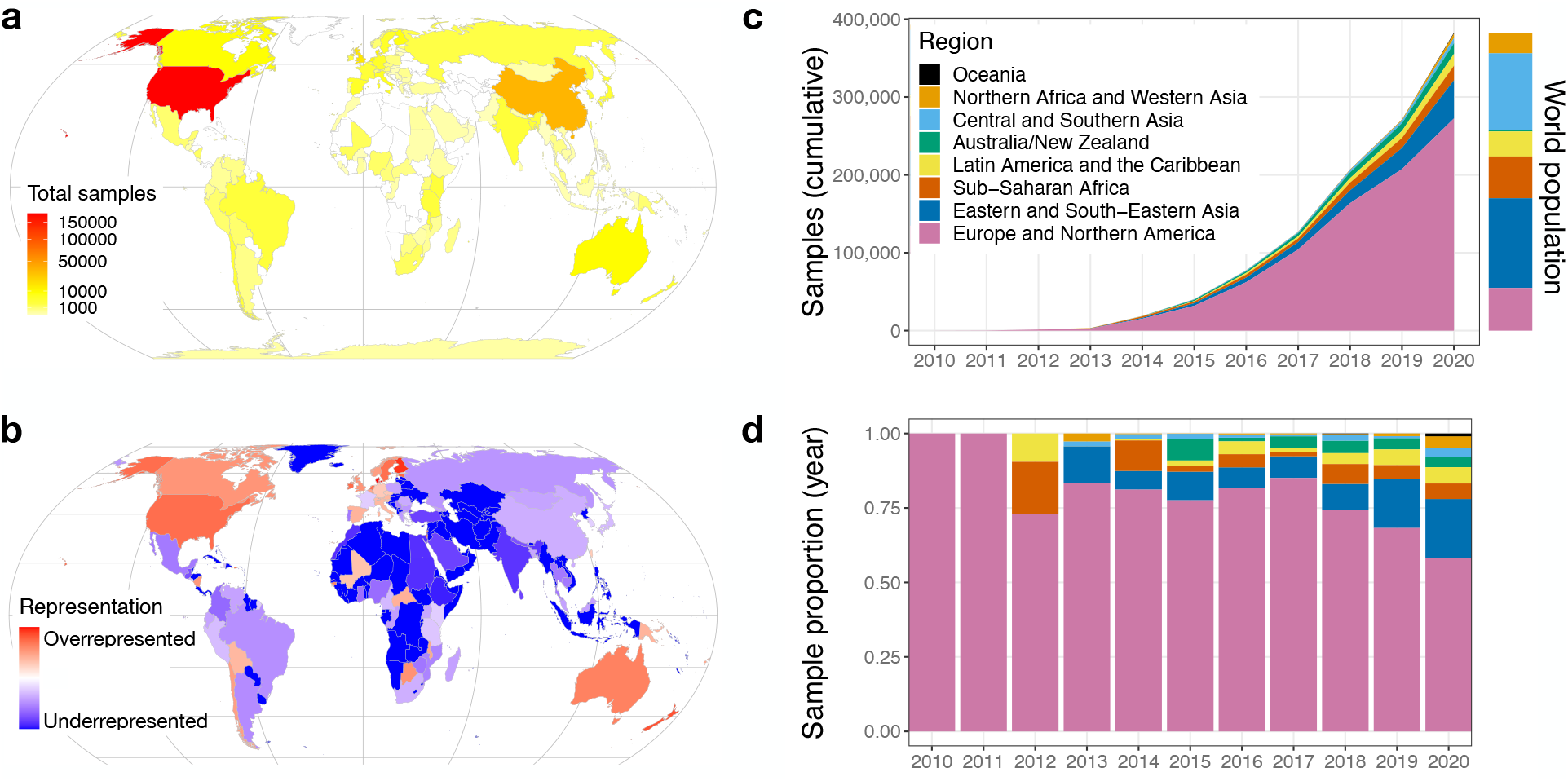
Global microbiome representation. (A) Total samples by country. The color of each country indicates the total number of samples originating in that country. (B) Relative representation by country. The color of each country indicates its representation in human microbiome datasets, relative to its share of world population. Red colors mark countries that are overrepresented relative to their population, and blue colors mark countries that are underrepresented. Countries with zero samples in the dataset are marked with dark blue. (C) Cumulative microbiome samples by world region. The x-axis indicates the year; the y-axis indicates the cumulative microbiome samples available at the end of that year. Colors indicate the cumulative microbiome samples from each of the world regions specified in the legend. The colored bar to the right of the plot indicates the share of the world population living in each of the regions using the same colors. (D) Proportion of annual samples. The x-axis indicates the year, and the y-axis indicates the proportion of samples from each world region published in that year. Colors correspond to the world regions shown in panel C.

To examine patterns of under- and overrepresentation of countries, we compared human microbiome sample counts to each country’s population, according to United Nations estimates for 2020 (“World Population Prospects 2019” 2019). The United States is dramatically overrepresented relative to its population: Though the country has about 4.3% of the global population, 40.2% of human microbiome samples originate there. Proportionally, Denmark is the most overrepresented country, with 11,497 samples from a country of about 5.8 million people (**Figure 1b**). Of the 235 countries and territories included in the United Nations population estimates, 120 have zero human microbiome samples available in these public databases.

To gain a better understanding of global representation in microbiome research, we grouped countries using the eight United Nations Sustainable Development Goals regions (“SDG Indicators” n.d.). We found that 71.2 percent of samples with a known location come from Europe and Northern America, a region that holds only 14.3 percent of the world’s population (**Table 2**). Proportionally, Australia/New Zealand has the most lopsided presence in the database: The region’s 30.3 million people is 0.4 percent of the population, but account for 3.1 percent of samples (**Figure 1c**). Central and Southern Asia is the most underrepresented region: It holds 25.8 percent of the population but makes up only 1.8 percent of microbiome samples. Northern Africa and Western Asia is the next-most underrepresented region, followed by Sub-Saharan Africa, which is home to 14.0% of the world’s population but is the source of 4.2% of human microbiome samples. These proportions indicate a person in Europe or Northern America is roughly 14 times more likely to be studied in a microbiome project than someone from Sub-Saharan Africa. The 47 countries on the United Nations list of “least developed countries” account for about 14 percent of the world’s population (“About LDCs” 2013), but 3.4 percent of microbiome samples; 29 of those countries have no samples at all (**Supplementary Table 2**). When we consider how global representation changed over time, we find that although samples from Europe and Northern America are over-represented, in recent years there is more representation for samples from other regions, most prominently eastern and south-eastern Asia (**Figure 1d**).

**Table 2.** Samples and population by region.

| **Region** | **Samples** | **2020**<br>**population**<br>**(estimated, in**<br>**thousands)** | **% of samples** | **% of samples**<br>**(known**<br>**location)** | **% of**<br>**population** | **Representation**<br>**proportion\*** |
| --- | --- | --- | --- | --- | --- | --- |
| Europe and Northern America | 272,544 | 1,116,506 | 61.3% | 71.2% | 14.3% | 4.97 |
| Eastern and South-Eastern<br>Asia | 49,007 | 2,346,709 | 11.0% | 12.8% | 30.1% | 0.43 |
| Sub-Saharan Africa | 18,651 | 1,094,366 | 4.2% | 4.9% | 14.0% | 0.35 |
| Latin America and the<br>Caribbean | 15,264 | 653,962 | 3.4% | 4.0% | 8.4% | 0.49 |
| Australia/New Zealand | 13,620 | 30,322 | 3.1% | 3.6% | 0.4% | 9.14 |
| Central and Southern Asia | 6,685 | 2,014,709 | 1.5% | 1.7% | 25.8% | 0.07 |
| Northern Africa and Western<br>Asia | 5,621 | 525,869 | 1.3% | 1.5% | 6.7% | 0.22 |
| Oceania | 1,178 | 12,356 | 0.3% | 0.3% | 0.2% | 1.94 |
| *Unknown* | 62,259 |  | 14.0% |  |  |  |
| Least developed countries | 15,254 | 1,057,438 | 3.4% | 4.0% | 13.6% | 0.29 |
| Rest of world | 367,457 | 6,737,361 | 82.6% | 96.0% | 86.4% | 1.11 |
| *Unknown* | 62,118 |  | 14.0% |  |  |  |

\*Representation proportion calculated by dividing a regions percentage of known samples by its percentage of population.

## Discussion

Our results show that the global distribution of human microbiome sampling is heavily skewed towards North American and European populations, both in total samples (**Figure 1a**) and in samples adjusted for population (**Figure 1b**). The United States is by far the greatest contributor to the database (**Table 1**), though this is slowly beginning to change as other countries’ contributions grow (**Figure 1d**). This neglect of most of the world’s population represents a problematic disparity in microbiome research. Since only a subset of the world’s populations are currently being studied, microbiome–disease associations identified may not hold in under-sampled populations (Gupta, Paul, and Dutta 2017; He et al. 2018). Additionally, by only sampling a subset of the global population, the diseases studied in the context of the microbiome are limited to diseases which impact that subset. To ensure greater global equity in the benefits of microbiome research, the field should prioritize and incentivize improved global representation of microbiome samples. Importantly, this approach should be grounded in benefitting the populations and communities sampled, rather than simply using these microbiomes as a tool to improve health in North American and European countries, as others have explained (Benezra 2020; Delgado and Baedke 2021).

There are several limitations to our study. Metadata quality is the primary hurdle in characterizing samples (Gonçalves and Musen 2019): It is possible that not all samples identified as human in this study are indeed from humans and could, for example, include studies using human gut microbiota transferred into mice. In addition, as it has been estimated that 20 percent of microbiome papers do not have publicly available data (Eckert et al. 2020), our study only examines the subset of microbiome studies that also shared their data in the largest international repositories. Our analysis also assumes that each sample represents a unique human individual and does not account for multiple samples originating from the same person. We have also limited our database search to three databases (Sequence Read Archive, DNA Data Bank of Japan, and European Nucleotide Archive); it is possible that different patterns of global representation are present in other databases, such as MG-RAST (Wilke et al. 2016) and gcMeta (Shi et al. 2019), though they are orders of magnitude smaller than the NCBI holdings.

To conclude, we analyzed the geographic origins of almost a half-million samples from the largest genomic repositories in the world. We find evidence that the human microbiome field may be repeating some of the same mistakes found in human genomics (Need and Goldstein 2009; Popejoy and Fullerton 2016), investing billions of dollars into research that excludes broad segments of the world. The field would benefit from a more global perspective on investigating the human microbiome’s relationship to health and disease.

## Materials and Methods

A list of samples was exported from the NCBI BioSample database (https://www.ncbi.nlm.nih.gov/biosample) between April and June 2021 using the search string “txid408170[Organism] AND biosample sra[filter] AND “public"[filter]", which requests all samples classified under the “human gut metagenome” taxon from the NCBI Taxonomy (https://www.ncbi.nlm.nih.gov/Taxonomy/Browser/wwwtax.cgi). The resulting sample IDs and all associated tags were loaded into a PostgreSQL database. We repeated this for all taxa described as human metagenomes (**Supplementary Table 3**). We note that the term “human gut metagenome” does not describe the sequencing technique used to generate the microbiome data, including shotgun metagenomics and amplicon sequencing -- specifically, 301,700 samples (72.0%) are associated with sequencing runs that list the library strategy as “AMPLICON."

We then looked in other NCBI taxa nested beneath “organismal metagenomes” that were not explicitly labeled “human” but were likely to contain some human samples (“Organismal Metagenomes” n.d.). We downloaded the metadata for samples classified under any NCBI taxa that was the “generic” version of a human one we’d already collected—the “blood metagenome” taxon is the generic version of the “human blood metagenome” taxon, for example. We downloaded all sample data for any generic taxa that had at least 1,000 samples (**Supplementary Table 3**), then evaluated the metadata to find which samples indicated they were taken from a human host. To do this, we used the value of the “host_taxid” field or, if that was blank, the value of “host,” to create a putative “host” value, and manually flagged any that explicitly indicated the sample was from a human—references to “human” or “homo sapiens,” for example, or if the host included words such as “patient” or “crew member” and did not indicate another species. We evaluated 4,395 unique “host” values for 173,038 samples and found 501 values assigned to 29,934 samples (17.3%) that indicated the host was a human. These were also included in the analysis.

We then used the NCBI eUtils API to find “runs” associated with each sample, so we could ensure all the BioSamples were associated with actual sequencing data. In the NCBI system, “runs” are the entities associated with sequencing data. We also used this API to obtain information on publication date, library strategy and the dates on which samples became publicly available. This resulted in a collection of 444,829 samples across 19 body sites (**Table 2**) after removing several hundred samples that were missing dates or sequencing data.

### Representation proportions

To determine which countries were over- or underrepresented relative to their populations, we obtained the 2020 population estimates for all countries as estimated by the United Nations (“World Population Prospects 2019” 2019). We used this to calculate two percentages for each country, one for the country’s share of the global population, and another for the country’s share of human microbiome samples. We then calculated a representation index: For countries with a higher sample percentage than population percentage, we divided the former by the latter to obtain a number indicating how many times more samples are present than expected. For countries with a lower sample percentage than population percentage, we took the negative reciprocal of this number, indicating (in negative numbers) the number one would have to multiply the sample count by to get the number that would be proportionally representative. The interim result leaves overrepresented countries with positive scores, and underrepresented countries with negative scores. After removing the scores for countries with 50 or fewer samples, we scaled the positive scores to fall between 0 and 100, and separately scaled the negative scores to fall between 0 and −100. We then plotted these on the map using the “pseudo_log” transformation to add more variation in the color-coding for the countries with middling scores. For the regional calculations (Figure 1c–d), we used top-level classifications from the same United Nations document. Antarctica is not included in a region, so those samples were added to the “Unknown” category for region-level calculations.

### Visualization

All figures were made using R and the ggplot2 package (Wickham 2009). Maps use the Equal Earth projection (Šavrič, Patterson, and Jenny 2019) and the rnaturalearth R package (South 2017).

## Acknowledgements

We thank Casey S. Greene and the members of the Blekhman lab for helpful discussions, and the Minnesota Supercomputing Institute for providing computational resources. This work was supported by NIH grant R35GM128716 (to R.B.).

## Data availability

All data has been deposited at Zenodo.org and is available at https://doi.org/10.5281/zenodo.5351180. This repository also includes the code used for data collection, along with the code used to generate each plot.

## Supplementary material

Supplementary data files and documentation are available at https://doi.org/10.5281/zenodo.5351180.

**Supplementary Table 1. Samples per tag.** Each row represents a single metadata field available for BioSample entries. The “samples” column indicates how many samples have a value for that field.

**Supplementary Table 2. Country-level data.** Each row represents a single country or territory as defined by the United Nations. There are 10 columns; see the supplementary documentation for a description of them.

**Supplementary Table 3. Samples by NCBI taxon.** Each row indicates a single entry in the NCBI Taxonomy Browser (https://www.ncbi.nlm.nih.gov/Taxonomy/Browser/wwwtax.cgi) related to the human microbiome. The “count” column indicates the total number of samples within each taxon.

